# MRSCloud: a Cloud-based MR Spectroscopy Tool for Basis Set Simulation

**DOI:** 10.1101/2022.03.22.485310

**Authors:** Steve C.N. Hui, Muhammad G. Saleh, Helge J. Zöllner, Georg Oeltzschner, Hongli Fan, Yue Li, Yulu Song, Hangyi Jiang, Jamie Near, Hanzhang Lu, Susumu Mori, Richard A. E. Edden

## Abstract

**Background:** Accurate quantification of in vivo proton magnetic resonance spectra involves modeling with a linear combination of known metabolite basis functions. Basis sets can be generated by numerical simulation using the quantum mechanical density-matrix formalism. Accurate simulations for a basis set require correct sequence timings, and pulse shapes and durations.

**Purpose:** To present a cloud-based spectral simulation tool ‘MRSCloud’. It allows community users of MRS to simulate a vendor- and sequence-specific basis set online in a convenient and time-efficient manner. This tool can simulate basis sets for 3 major MR scanner vendors (GE, Philips, Siemens), including conventional acquisitions and spectral editing schemes (MEGA, HERMES, HERCULES) with PRESS and semi-LASER localization.

**Study Type:** Prospective.

**Specimen:** N/A

**Field Strength/Sequence:** Simulations of 3T basis sets for conventional and spectral-editing sequences (MEGA, HERMES, HERCULES) with PRESS and sLASER localizations.

**Assessment:** Simulated metabolite basis functions generated by MRSCloud are compared to those generated by FID-A and MARSS, and a phantom-acquired basis-set from LCModel.

**Statistical Tests:** Intraclass correlation coefficients (ICC) were calculated to measure the agreement between individual metabolite basis functions generated using different packages. Statistical analysis was performed using R in RStudio.

**Results:** Simulation time for a full basis set is approximately 1 hour. ICCs between MRSCloud and FID-A were at least 0.98 and ICCs between MRSCloud and MARSS were at least 0.96. ICCs between simulated MRSCloud basis spectra and acquired LCModel basis spectra were lowest for Gln at 0.68 and highest for NAA at 0.96.

**Data Conclusion:** Substantial reductions in runtime have been achieved by implementing the 1D projection method, coherence-order filtering, and pre-calculation of propagators. High ICC values indicated that the accelerating features are running correctly and produce comparable and accurate basis sets. The generated basis set has been successfully used with LCModel.

## Introduction

Magnetic resonance spectroscopy (MRS) is a non-invasive method that uses an MRI scanner to detect radiofrequency (RF) signals from endogenous brain metabolites. Quantitative MRS has a broad range of preclinical and clinical research applications. Changes in metabolite levels can be observed in neurodegenerative, psychiatric and neurodevelopmental disorders (1,2) and are potential biomarkers for brain tumor phenotyping and tracking treatment response (3,4). A key challenge in MRS has been bridging from detection to quantification, i.e., access to robust, reproducible, and transparent analysis workflows that convert the acquired spectrum into measured metabolite levels. Although the signals derived from each metabolite change linearly with concentration, resolution between the signals of the large number of metabolites that are present in brain tissue is very limited. Inferring the concentration of individual metabolites from the complex mixture of signals that make up the in vivo spectrum is therefore challenging.

The most widely accepted approach to analyze MR spectra is linear combination modeling (LCM). In LCM, the acquired spectrum is modeled as a weighted sum of metabolite basis functions and other terms to describe background signals from mobile lipids and macromolecules and the spectral baseline. Metabolite basis sets can either be generated by numerical simulation or experimental acquisition of spectra from a set of single-metabolite phantoms. The latter is time-consuming, expensive and extremely demanding technically. Simulations based on the quantum mechanical density-matrix formalism (5) have become easier over time, with increased computing power and broadened access to simulation tools (6–12). Accurate simulated basis sets can only be generated using accurate acquisition parameters, including experimentally applied vendor-appropriate RF pulse shapes, correct timings and sufficient spatial resolution (13,14). Although several simulation tools are available (6–12), their use involves a significant learning curve. Only a minority of expert MRS groups posses the detailed sequence knowledge to implement accurate basis-set simulations locally. Most LCM analyses rely upon basis sets generated off-site, often with approximate, rather than exact-match parameters. This approach is feasible for fitting the large, relatively well-separated singlet resonances of NAA, choline and creatine in short-TE spectra. However, using approximated basis sets is less tolerable for lower-concentration coupled spin systems and for more complex (i.e. longer-TE and edited) experiments. Two substantial shifts have recently occurred in MRS analysis. The predominant software product in the field ‘LCModel’ (15) is no longer actively developed or supported, and at the same time a new generation of fuller-function, often open-source analysis software has been released, including FSL-MRS (16), spant (17), and Osprey (18). These tools perform LCM but access to appropriate basis-sets persists as a hurdle in broadening access to the state-of-the-art MRS analyses.

The purpose of this manuscript is to introduce MRSCloud, a cloud-based basis-set simulation tool. MRSCloud allows users to generate density-matrix-simulated basis sets through a simple web interface. It builds upon FID-A simulation functionality (6). FID-A can perform spatially resolved simulations using shaped pulses, but these are not optimized for simulation speed. Density-matrix simulations can be substantially accelerated by using the one-dimensional (1D) projection method (19) and applying coherence pathway filters (20,21) to replace phase cycling. Cloud-based basis-set generation reduces the expertise required to carry out spectral analysis and quantification. Users require basis sets for conventional and edited MRS experiments (22), including Hadamard-encoded editing (23,24). MRSCloud implements the 1D projection method (19), coherence pathway filters (21) and pre-calculation of propagators within the FID-A code base. Parameters specified by the user include localization method (PRESS (25) or semi-LASER (26,27)), vendor (GE, Philips, Siemens and the standardized universal sequence (28)), spectral editing schemes (unedited, MEGA (29), HERMES (23), HERCULES (24)), metabolite list, spatial resolution, echo time (TE) and editing pulse frequencies. It allows community users to generate vendor-, sequence-, and editing-experimental-specific basis sets that are appropriate for their studies.

## Methods

### Development of Simulations

This cloud implementation builds upon the simulation tools of FID-A, implementing three accelerations, which are described in the following sections.

### 1D projection method

Spatially resolved simulations are time-consuming, but necessary in order to capture spatial heterogeneity of coupling evolution. In the original FID-A implementation of spatial simulations, each spatial direction was governed by a loop, and the full sequence was simulated for each spatial location. Simulating *n_x_*, *n_y_*, and *n_z_* spatial points in the x-, y- and z-direction, respectively amounts to performing *n_x_* × *n_y_* × *n_z_* simulations of the sequence. Zhang et al. (19) demonstrated that a more efficient way to achieve three-dimensional simulations is to average across each spatial dimension after simulating each one-dimensional frequency-selective RF pulse. In PRESS localization, for example, the first spatial averaging can be performed after simulation of the slice-selective excitation pulse (and the appropriate accompanying rephase gradient). In the MRSCloud implementation, the excitation pulse is not explicitly simulated, but approximated by an ideal excitation. An additional spatial averaging step is performed after the first slice-selective refocusing pulses, and then finally after the second slice-selective refocusing pulse. The total simulation time decreases from the order of (*n_x_* × *n_y_* × *n_z_*) to (*n_x_* + *n_y_* + *n_z_*)/3.

### Coherence pathway filters

Slice-selective refocusing relies upon coherence pathway selection to suppress signals from outside the intended slice. In the sequence itself, this is usually achieved both by phase cycling (21) and the coherence transfer pathway selection gradient scheme. Typically, simulations are not performed with sufficient spatial resolution for simulated gradients (7) alone to accurately suppress out-of-voxel signal, and simulations rely upon phase cycling. The commonly used EXORCYCLE (30) scheme is a suitable four-step phase cycle; independent EXORCYCLE of the two refocusing pulses results in a sixteen-step phase cycle (i.e. the simulation is repeated 16 times with different RF pulse phases and simulated receiver phase). In FID-A, it is common to approximate this with a half-EXOCYCLE on each pulse, resulting in a four-step phase cycle.

The MRSCloud implementation removes the need for phase cycling by applying coherence pathway filters (7,9,10). The different elements of the density matrix can each be classified in terms of their coherence order, and unwanted elements can be zeroed (equivalent to perfect coherence transfer pathway selection). Each simulated RF pulse in the sequence is immediately followed by a coherence-order filter, so that only density-matrix elements of the intended order are propagated further in the simulation.

### Pre-calculation of propagators

Within FID-A, shaped pulses are calculated elementwise each time they are applied. For example, a 256-point shaped refocusing pulse is simulated as 256 individual 3D rotations. This involves a substantial amount of redundant calculation that can be avoided by pre-calculation of the pulse propagators. Simulating each pulse then becomes a matter of applying the appropriate propagator, rather than serially calculating and then applying 256 individual propagators. The saving arises both because refocusing and editing pulses are repeated within the sequence and, in cases where the previous two accelerations have not been implemented, for phase cycling and spatial purposes.

Figure 1a illustrates the accelerated MRSCloud simulation for the example of MEGA-PRESS, in terms of its constituent propagation operations.

**Figure 1:**
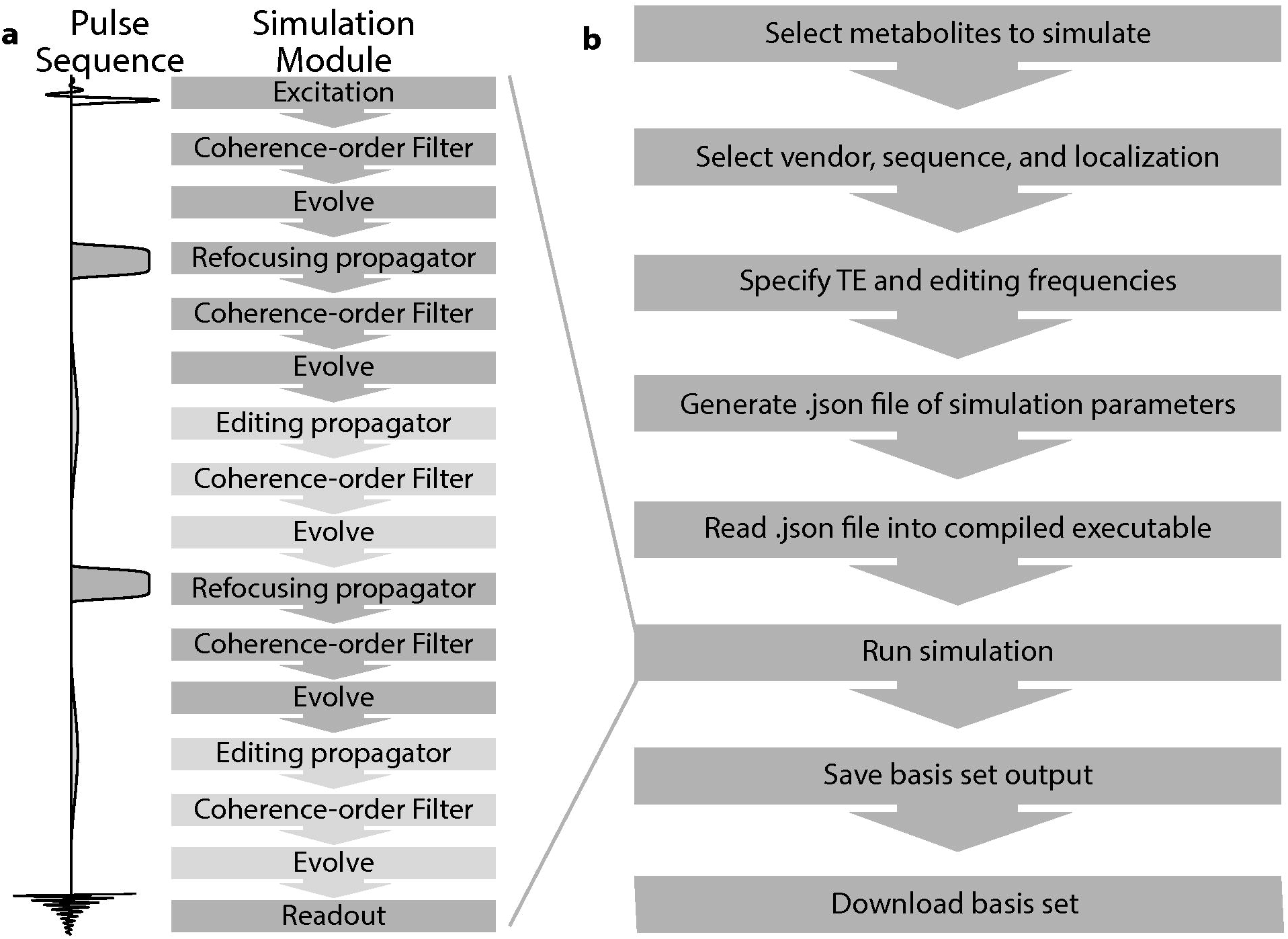
Flowchart for a) simulation module along the sequence diagram and b) MRSCloud pipeline

### GUI specification of simulation parameters

Figure 1b outlines the pipeline for a given MRSCloud simulation. Firstly, the user specifies the metabolites to be simulated, from a list of 32. Of those, 25 use FID-A spin-system definitions (6), many of which are parameterized from (31,32), augmented by other metabolites including cystathionine, ethanolamine, homocarnosine, lysine, phosphorylethanolamine, threonine and valine (31,33,34). Next the user specifies the vendor (GE, Philips, Siemens or Universal), sequence options (Unedited, MEGA, HERMES or HERCULES (23,24,29)) and localization method (PRESS or semi-LASER). TE can be defined for Unedited and MEGA simulations, and is fixed at 80 ms for HERMES and HERCULES. Editing frequencies of ON and OFF scans are user-defined for MEGA (default edit-on/-off: 1.9/7.5 ppm) and fixed for HERCULES (1.9/4.18/4.58 ppm) and HERMES (GABA/GSH 1.9/4.56 ppm). The GUI for MRSCloud is shown in Figure 2. Once the user presses submit, user inputs are stored in a declaration file (.json) that is loaded into the precompiled simulation executable to execute the desired simulation. The basis set output is then saved on the server for user download. All code was written in MATLAB (R2020b, MathWorks, Natick, USA), and compiled as an executable file to run on the server with MATLAB Runtime. Simulations are performed with a simulated B_0_ field strength of 3.0 T for Philips and GE, and 2.89 T for Siemens.

**Figure 2:**
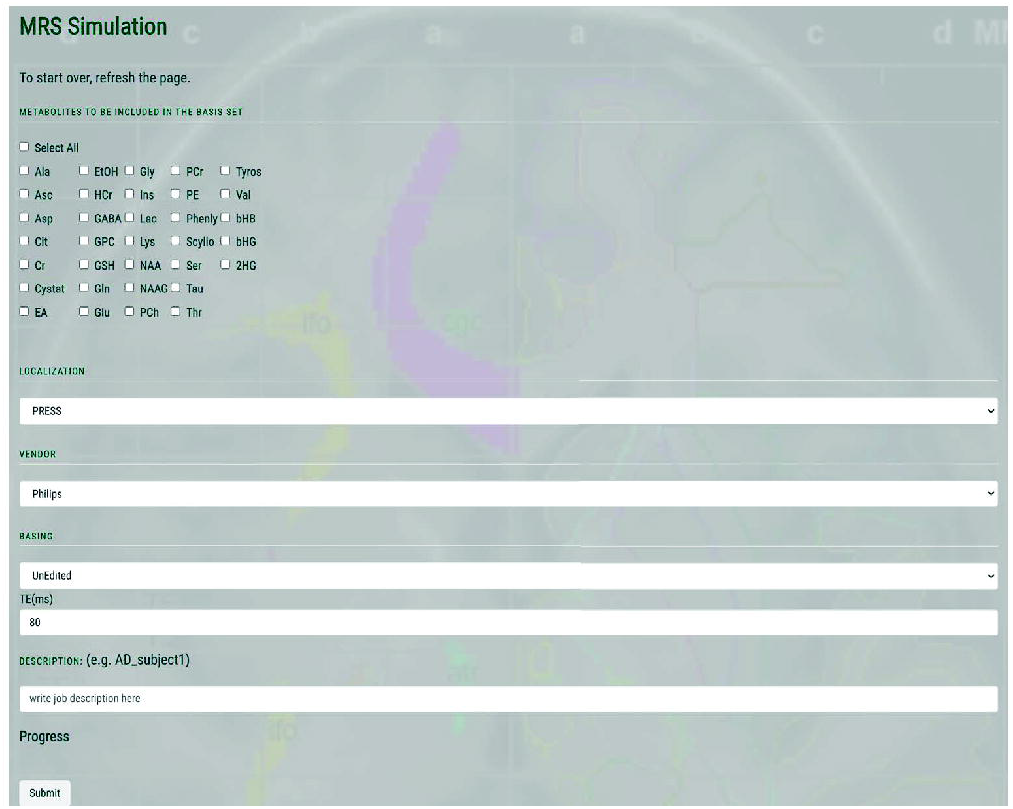
GUI of MRSCloud

### RF waveforms

MRSCloud simulations allow exact simulation of vendor-specific RF pulse shapes e.g. (28,35,36). For sLASER, GOIA-WURST refocusing pulses (bandwidth 8 kHz, duration 4.5 ms, B_1_ 15 uT) are simulated (37,38). Editing pulse durations for MEGA sequences are determined from TE (14 ms for TE = 68 ms; 20 ms for TE = 80 ms). Editing pulses were generated using a sinc-Gaussian pulse for Philips and GE (bandwidth 88 Hz, duration 14 ms), and the standardized universal sequence (bandwidth 86 Hz, duration 14 ms) and Hanning-filtered pulse (bandwidth 48 Hz, duration 25.6 ms) for Siemens. 20-ms (bandwidth 62 Hz) editing pulses are simulated for HERMES and HERCULES; dual-band editing pulses are generated using a cosine modulation of the editing pulse.

### Spatial Resolution

A spatial array of simulations (of 101×101 resolution, by default) is carried out across a field of view that extends 50% larger than the nominal voxel size in the two dimensions defined by refocusing pulses. In order to investigate the impact of spatial resolution, MRSCloud simulations were performed with resolutions of 21×21, 41×41 and 101×101. Edited GABA signals from MEGA, HERMES and HERCULES for PRESS and sLASER at TE of 80 ms were compared using intraclass correlation coefficient (ICC).

### Testing and validation

The speed of simulation was tested locally on a MacBook Pro (2.3 GHz Quad-Core Intel Core i5, 16 GB LPDDR3 memory) and on the MRSCloud server. Simulations were carried out using PRESS localization with un-edited, MEGA, HERMES and HERCULES sequences. 25 major metabolites (Ala, Asc, Asp, Cit, Cr, EtOH, GABA, GPC, GSH, Gln, Glu, Gly, mI, Lac, NAA, NAAG, PCh, PCr, Phenyl, Scyllo, Ser, Tau, Tyros, bHB and 2HG) were included. For local testing, basis sets were generated using 21×21, 41×41 and 101×101 spatial points. For MRSCloud server testing, simulations were performed using the default setting of 101×101 spatial points. As server performance is usage-dependent, simulation speed was tested at three different times of day and the average simulation time was reported.

The first stage of validation of the MRSCloud basis sets is to compare them to equivalent basis sets produced by the release version of FID-A (i.e. without the implementation of coherence pathway filtering and pre-calculation of propagators). As downloaded in May 2021, FID-A includes the 1D projection method, but not the other improvements implemented in MRSCloud. Basis functions were simulated in MRSCloud and FID-A for 8 commonly used metabolites (Cr, GABA, Gln, Glu, mI, Lac, NAA, PCh) with PRESS localization at TEs of 30 ms and 68 ms. Simulations were performed locally in MATLAB using the pre-compiled MRSCloud code and the official FID-A package version. Spectral results obtained from the two methods were compared using ICC. Discrepancies between basis sets were quantified using a normalized rootmean-square value. In the second stage of validation, an MRSCloud basis set (un-edited Philips PRESS TE of 30 ms) was compared to an equivalent one generated with the MARSS software (available online: http://juchem.bme.columbia.edu/mr-spectroscopy-basis-sets). Again, ICC and normalized root-mean-square differences were calculated for each metabolite. Finally, an MRSCloud simulated basis set was compared to an experimentally acquired basis set, available for LCModel (press_te30_3t_01a.basis; http://.s-provencher.com/lcm-basis.shtml). All statistical analyses were performed using R (Version 4.0.2) in RStudio (Version 1.2.5019, Integrated Development for R. RStudio, PBC, Boston, MA).

## Results

MRSCloud has been successfully established, offering web-based basis-set simulation for the community through the portal: https://braingps.anatomyworks.org/mrs-cloud. Basis function profiles for MEGA-PRESS (TE = 68 ms) are shown in Figure 3.

**Figure 3:**
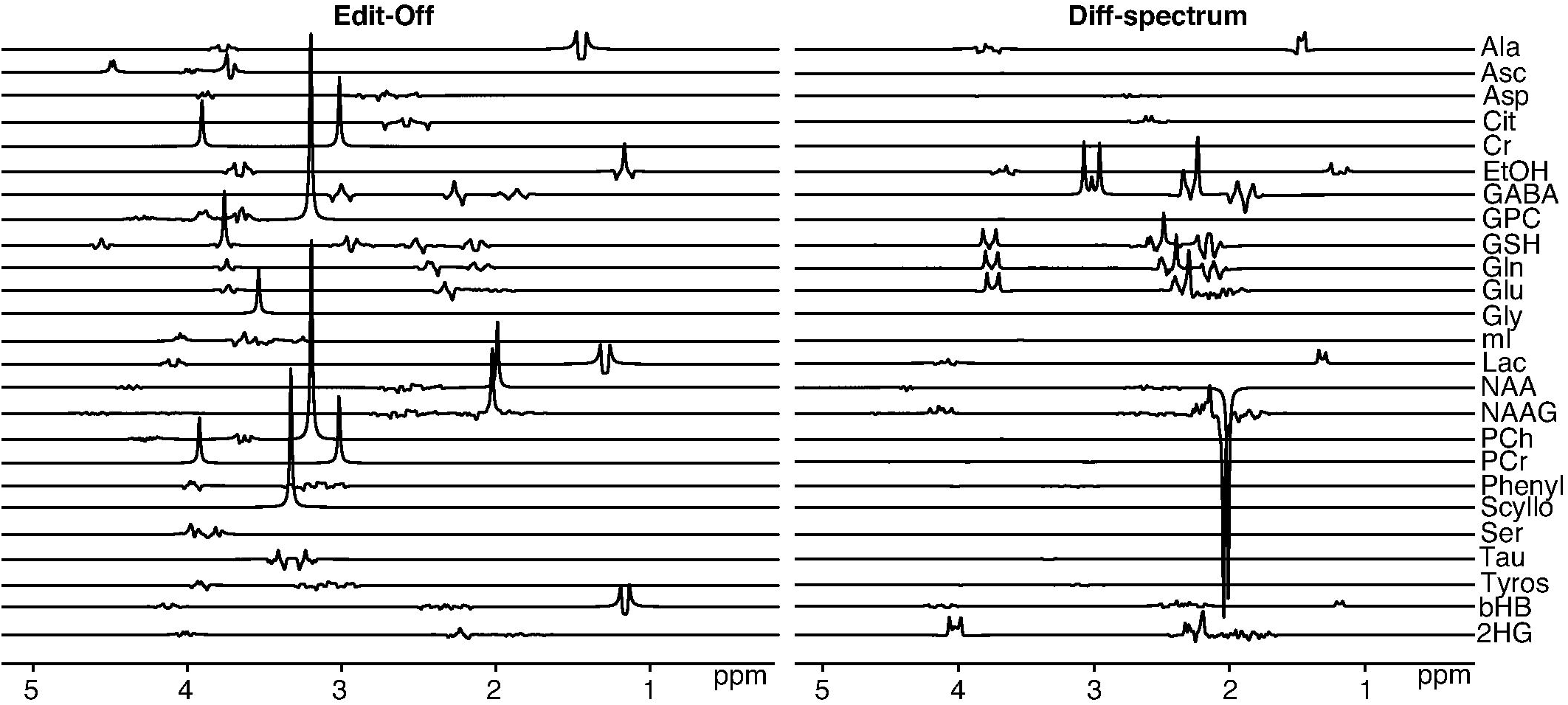
Basis set components for MEGA-PRESS GABA-editing at TE 68 ms simulated using MRSCloud

MEGA-PRESS, HERMES and HERCULES simulations of the GABA-edited difference signal at spatial resolutions of 21×21, 41×41 and 101×101 are shown in Figure 4a for PRESS and 4b for sLASER localization. GABA integrals at 3 ppm were slightly larger for larger numbers of spatial points. Simulation time of 101×101 spatial points for GABA, a strongly coupled 6-spin system, was less than 1 minute for all three sequences locally. ICCs among the different spatial resolutions were higher than 0.99 for all three sequences in both PRESS and sLASER.

**Figure 4:**
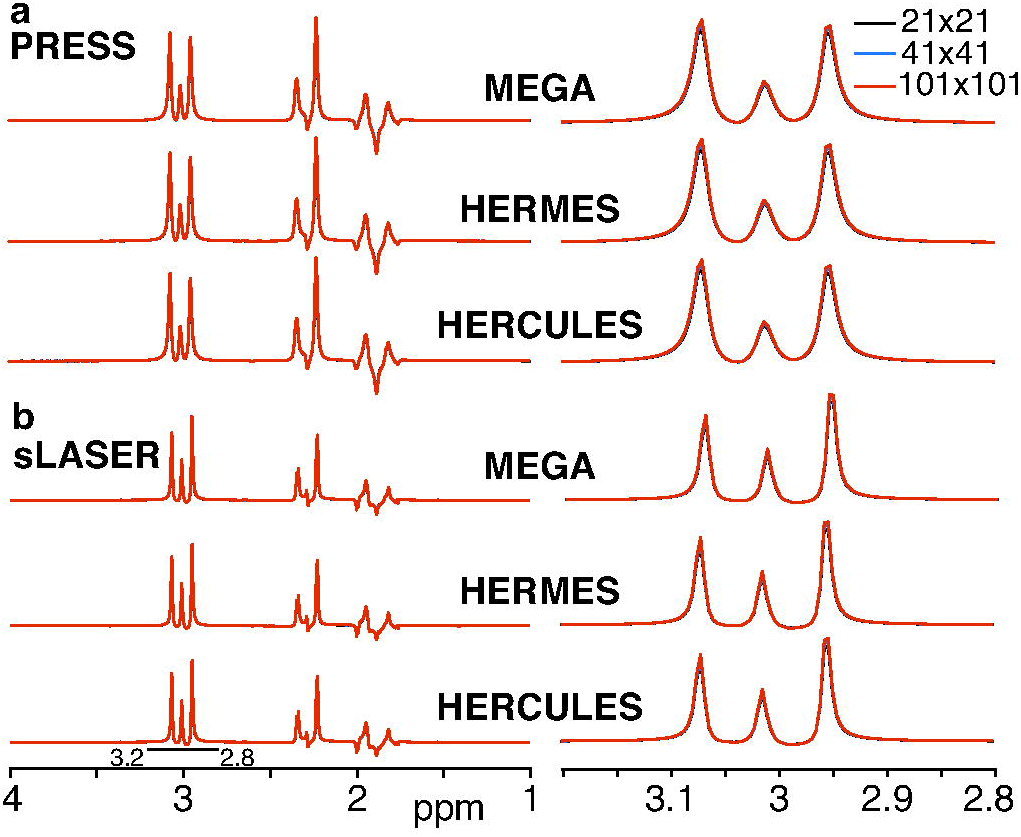
Simulated GABA-edited difference spectra using MEGA, HERMES and HERCULES with a) PRESS and b) sLASER localization, at 21×21, 41×41 and 101×101 spatial resolutions. The 3-ppm edited signal is expanded on the right.

Time for simulations is reported in Table 1. Simulation times increase with the number of spatial points and the complexity of the sequence. The local simulation time was between 3 and 12 minutes for a full basis set. Simulation time on the server was longer between 11 and 25 minutes for different sequences. Time required to simulate a full basis set on the server is similar at different times.

**Table 1:** Time for simulating a full basis set (25 metabolites) for different sequences using Philips PRESS localization.

|  | TE | Local Simulation Time (sec) |  |  | Server (sec) |
| --- | --- | --- | --- | --- | --- |
| | | $21^2$ | $41^2$ | $101^2$ | $101^2$ |
| Un-edited | 30 | 183 | 215 | 270 | 676 |
| MEGA | 68 | 300 | 320 | 388 | 947 |
| HERMES | 80 | 545 | 628 | 719 | 1444 |
| HERCULES | 80 | 556 | 651 | 732 | 1490 |

Un-edited PRESS simulations at TE = 30 ms and TE = 68 ms using MRSCloud were comparable to FID-A simulations, as shown in Figure 5. Local simulation time for MRSCloud was reduced by a factor of 9 compared to the native FID-A simulation. Results from all simulated metabolites indicated both methods generated very similar lineshapes and signal patterns. ICCs were higher than 0.98 for all tested metabolites, indicating high similarity for the basis functions generated using MRSCloud and the native FID-A simulations. Discrepancy for the corresponding metabolites between the two methods is presented in the same figure. The normalized root-meansquare value of the differences are under 1% for all metabolites, as shown in Table 2.

**Figure 5:**
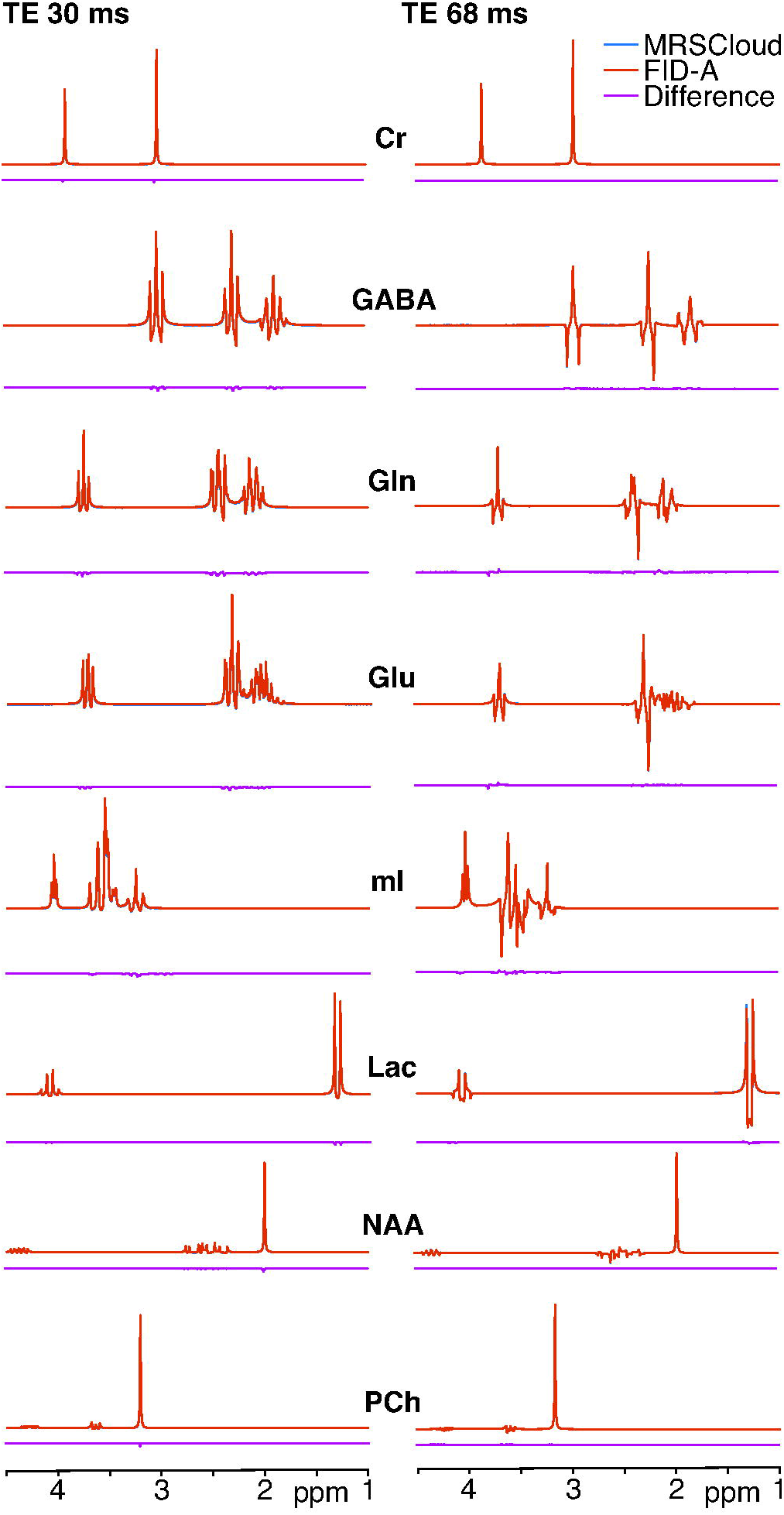
Comparison between MRSCloud and local FID-A short-TE basis functions for 8 major metabolites (41×41 spatial points).

**Table 2.** Normalized root-mean-square differences between MRSCloud basis spectra and FID-A, MARSS and LCModel.

| Compared to: | Cr | GABA | Gln | Glu | mI | Lac | NAA | PCh |
| --- | --- | --- | --- | --- | --- | --- | --- | --- |
| FID-A TE 30 | 0.088% | 0.3% | 0.38% | 0.59% | 0.28% | 0.1% | 0.098% | 0.06% |
| FID-A TE 68 | 0.0031% | 0.054% | 0.023% | 0.046% | 0.021% | 0.17% | 0.0073% | 0.0022% |
| MARSS | 3.8% | 6.0% | 5.6% | 8.3% | 5.8% | 3.9% | 3.3% | 3.2% |
| LCModel | 6.2% | 11.3% | 22.5% | 11.6% | 4.7% | 9.6% | 2.2% | 2.9% |

Results from MRSCloud agreed well with those from MARSS as shown in Figure 6. Differences were more pronounced for coupled spin systems like GABA, Glu and Gln. ICCs were higher than 0.96 for all tested metabolites. The normalized root-mean-square differences are below 9% for all metabolites, and shown in Table 2.

**Figure 6:**
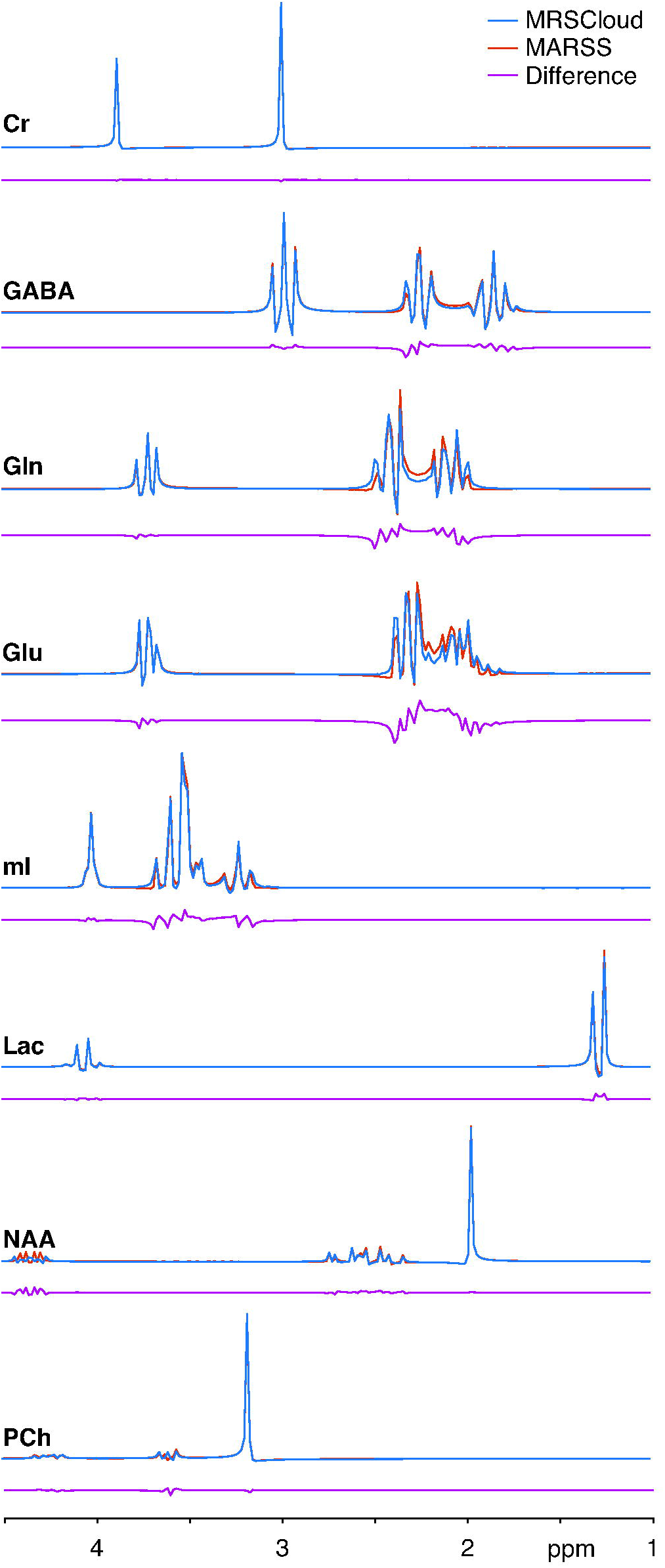
Comparison between MRSCloud and MARSS in 8 major metabolites for PRESS at TE 30 ms.

The MRSCloud-simulated spectra were in reasonable agreement with the experimentally measured basis set that is available from the LCModel as shown in Figure 7. ICCs were lowest for Gln at 0.68 and highest for NAA at 0.96 which corresponded to the largest and smallest rootmean-square values for LCModel in Table 2.

**Figure 7:**
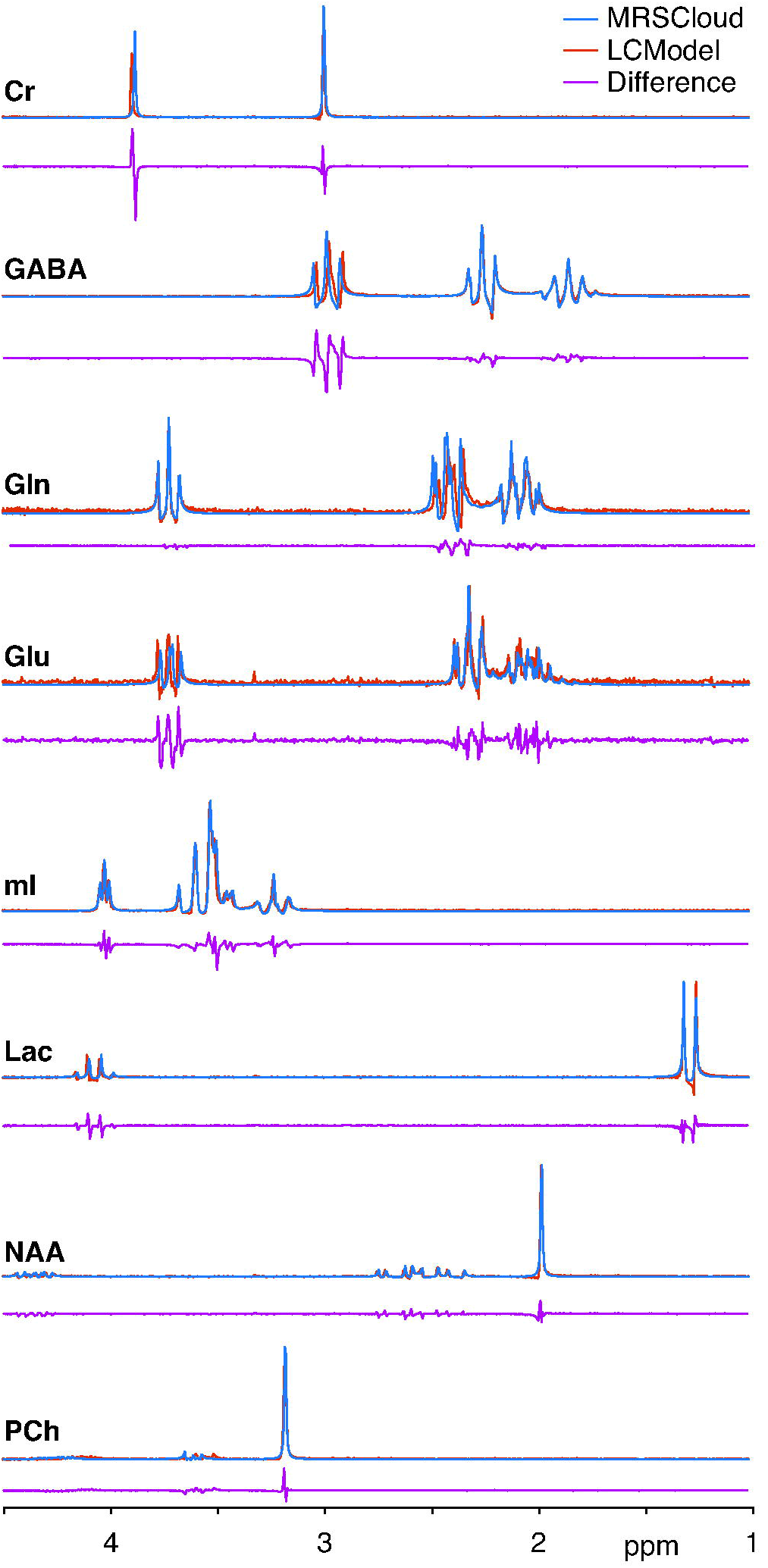
Spectral shape comparison between MRSCloud and LCModel

## Discussion

MRSCloud is a cloud-based simulation tool for the MRS community, offering fast and reliable basis set generation for up to 32 metabolites. This improves access to basis sets that more closely represent the timing and RF pulse shapes of vendor specific sequences. Substantial reductions in runtime have been achieved by implementing the 1D projection method, coherence-order filtering, and pre-calculation of propagators.

Each implementation benefits the speed and precision of the simulations. The 1D projection method accelerates the process by decreasing the number of simulations to the factor equivalent to the square of the number of spatial points. This implementation allows efficient generation of metabolite responses with high spatial resolution, which is necessary to accurately reflect the spatial effects of imperfect excitation and refocusing, i.e. realistic slice profiles. Coherence order filtering helps the selection of appropriate coherence pathway to avoid the necessity to simulate full phase cycling, which accelerates the processing time by a factor equivalent to the number of phase cycles. This implementation also improves quality of simulations by providing the equivalent effect of including gradient to handle unwanted magnetizations or coherences. It is important especially for simulation using PRESS localization and strongly coupled matebolites sensitive to chemical shift displacement error (39). Pre-calculation of the propagator has been used to replace the multiple calculations of the density matrix time evolution operator for each individual pulse. It reduces the number of calculations from the square of the number of pulse samples to a 1-time calculation. All 3 implementations dramatically reduce the simulation time. Table 1 shows that a full basis set simulation takes approximately 11-25 minutes on MRSCloud (with negligible variation over the course of a day) and 3-12 minutes locally for different sequences. The time difference between the local machine and the cloud server is partly due to the performance of the processors and the older version of MATLAB being used for the server to run the compiled executable. According to the release note, the parallel computing toolbox in MATLAB has been upgraded since 2013-2014. The old version of parallel computing features such as the *matlabpool* have been succeed by the current *parpool* functions.

Simulations of edited sequences with different spatial resolutions (21×21, 41×41 and 101×101) have been carried out using MRSCloud. Differences are small as indicated by the high ICC values. Our result supports a similar comparison in another study for the spectral shape of unedited GABA (9). Coherence order filtering is particularly important because it accurately modeled the dephasing effects of crusher gradients especially when the number of spatial points is small. Although MRSCloud is developed based on FID-A, comparison between both packages is necessary as MRSCloud has implemented multiple accelerating strategies. Potential discrepancies in lineshape could be driven by the coherence transfer pathway selection offered by direct filtering of the density matrix, as compared to incomplete phase-cycling of refocusing pulses. However, both ICCs and root-mean-square values indicated that the lineshapes and signal patterns for all compared metabolites match very well, which verified that the coherence order filtering produced results equivalent to the traditional phase cycle method, in a much more time-efficient manner. MRSCloud has also been compared with MARSS. Similar spectral shape and small differences from basis functions, as shown in figure 6, suggest that basis sets from both packages are comparable. High ICCs indicated good agreement between simulated spectra from the two methods. More complex coupled metabolites such as GABA, Gln and Glu have a greater discrepancy of root-mean-square values (5-8%). Such differences are possibly caused by the use of different waveforms and timings which affect more significantly to the multiplet coupled metabolites after signal evolution. LCModel has been the most representative software for spectral modeling and analysis for over two decades. However, basis set comparison between MRSCloud and LCModel is challenging because limited information has been provided for the acquisition of the basis sets. Information from the user’s manual and observation from the vendor provided basis set namely ‘press_te30_3t_01a.basis’ in LCModel suggested that it is measured using phantom rather than simulation. Concentration, composition and temperature of the phantom could be the leading factors for discrepancy between measured and simulated results. Waveforms and timings for the measured basis set are not well documented which are likely to be different from the MRSCloud simulation. All those factors contribute to the high ICC values especially for the more complex coupled metabolites including GABA, Gln and Glu. However, basis set generated using MRSCloud has been successfully used in LCModel.

Recently, MEGA has become a product sequence in the latest releases in at least two of three main vendors. However, parameters including pulse shape and duration, bandwidth and maximum B1 power of the slice-selective refocusing RF pulses have not standardized. Also, in Siemens, frequency of the spinning hydrogen protons is slightly different due to its field strength at 2.89 T for a ‘3 T’ scanner. Those differences from vendors have been taken into account for simulation. MRSCloud users can be benefited from the flexibility on selecting edited target at different frequency for MEGA which allow them to obtain basis set designed for specific metabolites such as ethanol (1.17 ppm/3.65 ppm) and phosphorylethanolamine (3.22 ppm/3.98 ppm). Similar benefits can be experienced from the flexibility of TE which can be used to optimize the refocusing of specific metabolites such as GABA (68 ms) and Lac (140 ms). For more advanced editing sequences such as HERMES and HERCULES, pulse shapes and durations of editing pulses and editing frequency have been considered for different vendors and predefined for users. The internal setting for HERMES is for editing GABA/GSH (1.9 ppm/4.56 ppm). Other metabolites of interest such as NAA/NAAG (4.38 ppm/4.62 ppm) would be available in a later update. In MRSCloud, there is an option of ‘Universal’ beside the three major vendors. It is dedicated to simulating basis sets for users of the universal sequence (28). Although it is not a vendor generic sequence, it has standardized the pulse shape and duration, bandwidth, maximum B1 power and other parameters including TE1, TE2 for multi-vendors. Basis set for the universal sequence can be generated using MRSCloud which would facilitate the promotion of using this sequence.

Not all pulse shapes for different sequences are available from vendors. Although duration and bandwidth of the slice-selective refocusing RF pulses can be found in literature, the exact waveform is unknown for specific vendor. Furthermore, there is no standardized setup for MEGA PRESS. Although the two editing pulses are always placed in the middle between the excitation and the second refocusing pulse, and the second refocusing pulse to the signal acquisition, the first refocusing pulse can be located differently across vendors or in different version of software releases based on different gradient settings. Hence, TE1 and TE2 values are based on current release of software. Similar for sLASER, our setup is based on the available information obtained from the software release. However, each vendor may have different setup on the gradient scheme, sweep width and slew rate etc. The Minnesota group has published a standardized across-vendor sLASER scheme (38). However, it is a new scheme and has not been fully adopted by all major vendors. Implementation of a universal standardized PRESS and sLASER would be ideal for simulation because parameters are fixed and accurate. Before we can achieve that, simulations based on vendor and sequence using dedicated parameters are needed for proper spectral modeling. Another possible addition to MRSCloud is STEAM. Even though it is not as commonly used as the other localizations mentioned previously at clinical field strengths, it is still worth to be included in the future since it can be used in low TE spectral acquisition for specific study. Other functions including simulation of a specific moiety in interested metabolites could be included.

In conclusion, MRSCloud facilitates the generation of basis set for the MRS community in a convenient and time-efficient style. It provides simulation of the common metabolites in addition to some specific ones related to brain pathology such as tumor. Basis sets are saved in different formats that are compatible with commonly used modeling packages. The purposed work hoped to ease the prior knowledge required to simulation an accurate basis set so that spectral analysis can be carried out in clinical sites or academic units without the present of a spectroscopist.

## Acknowledgement

This work was supported by NIH grants R01 EB016089, R01 EB023963, R21 AG060245, R00 AG062230, K99 DA051315 and P41 EB031771.

